# A low-cost mouse cage warming system provides improved intra-ischemic and post-ischemic body temperature control – application for reducing variability in experimental stroke studies

**DOI:** 10.1101/2020.12.30.424355

**Authors:** Sung-Ha Hong, Jeong-Ho Hong, Matthew T. Lahey, Liang Zhu, Jessica M. Stephenson, Sean P. Marrelli

**Affiliations:** Department of Neurology, McGovern Medical School, the University of Texas Health Science Center, Houston, TX USA; Department of Neurology, Brain Research Institute, Keimyung University School of Medicine, Dongsan Medical Center, Daegu, South Korea; Department of Medicine, McGovern Medical School, the University of Texas Health Science Center, Houston, TX USA

**Author notes:** For correspondence: Dr. Sean P. Marrelli, McGovern Medical School, 6431 Fannin Street, MSE R344, Houston, TX 77030 USA, Voice.

**Keywords:** brain and body temperature in stroke outcome, improved rigor in experimental stroke, mouse temperature regulation in stroke, post-stroke temperature management, temperature support cage

## Abstract

Experimental guidelines have been proposed to improve the rigor and reproducibility of experimental stroke studies in rodents. As brain temperature is a strong determinant of ischemic injury, tight management of brain or body temperature (Tcore) during the experimental protocol is highly recommended. However, little guidance is provided regarding how or for how long temperature support should be provided. We compared a commonly used heat support method (cage on heating pad) with a low-cost custom built warm ambient air cage (WAAC) system. Both heat support systems were evaluated for the middle cerebral artery occlusion (MCAo) model in mice. The WAAC system provided improved temperature control (more normothermic Tcore and less Tcore variation) during the intra-ischemic period (60 min) and post-ischemic period (3 hrs). Neurologic deficit score showed significantly less variance at post-stroke day 1 (PSD1) in WAAC system mice. Mean infarct volume was not statistically different by heat support system, however, standard deviation was 54% lower in the WAAC system group. In summary, we provide a simple low-cost heat support system that provides superior Tcore management in mice during the intra-ischemic and post-ischemic periods, which results in reduced variability of experimental outcomes.

**Highlights:**

- We describe the fabrication of a low-cost mouse cage warming system (warmed ambient air cage; WAAC system) that can be assembled and applied in any stroke laboratory.
- The WAAC system provides more precise control of post-stroke mouse body temperature compared with traditional heating pad warming system.
- The more precise control of post-stroke core temperature reduces variability in some experimental measures in more severely injured mice.

## Introduction

Many neuroprotective agents or strategies have shown significant benefit in experimental stroke models, only to fail to demonstrate similar benefit in clinical trials. As a result, an evolving set of experimental guidelines have been proposed by multiple investigators or groups in an effort to increase the rigor of stroke studies and thereby increase the probability for translation into effective clinical therapies(1999; Dirnagl, 2006; Fisher et al., 2009; Kahle and Bix, 2012; Percie du Sert et al., 2017). One significant component of these recommendations includes more rigorous monitoring and control of key physiological variables (including body and brain temperature) in stroke models. Because body/brain temperature is highly correlated with stroke outcome(Buchan and Pulsinelli, 1990; Cao et al., 2014; Cao et al., 2017; Liu and Yenari, 2007), variability in temperature (both during surgery *and* recovery) can lead to significant variability in outcomes. These effects can be especially pronounced in small rodents (rats and mice) due to their potential for rapid and significant temperature changes. However, current recommendations are vague or insufficient regarding temperature measurement or maintenance during the intra-ischemic or post-ischemic period. For example, it is recommended to provide feed-back controlled heat support during surgery, but temperature management during the awake portion of the occlusion or during the first few hours of reperfusion are not specifically addressed. Given the high dependence of ischemic injury on intra- and post-ischemic brain and body temperature, we suggest that more consistent measurement and tighter control of body temperature should be a prominent component of the overall strategy for reducing variability in experimental stroke studies.

The present study sought to evaluate the effect of tight temperature control during the full intra-ischemic period and the first 3 hours of reperfusion on stroke outcome variability. We developed a novel low-cost surgical recovery cage that provides more consistent and precise temperature support versus the conventional “cage on a heating pad” strategy. We found that housing mice in these recovery cages provided superior control of body temperature (measured by wireless transponder) and less variance in outcome measures compared with heating pad temperature support, particularly in mice with more severe stroke injury. Our results support the benefit of tighter body temperature control for reducing variability of stroke outcome measures. Furthermore, we present a simple low-cost cage warming device that can be easily incorporated into standard mouse holding cages, thus enabling the widespread use of this cage warming system.

## Materials and Methods

### Warmed Ambient Air Cage (WAAC) Design

The warmed ambient air cage (WAAC) was created by installing a regulated heating module on the wire rack of a ‘shoe box’ mouse cage. The heating module consisted of a thermostat, heater, and small circulating fan designed for temperature control of reptile habitats (IncuKit MINI, IncubatorWarehouse.com) (**Fig 1**). The fan was configured to direct air upward and then around the cage (i.e. not directly blowing on the mouse). However, as provided, the fan was too strong for the small mouse cage. We therefore modified the module by adding a 3D printed baffle to reduce the circulating air flow (**Fig 1B**). The effect of the added baffle was evaluated by measuring the clearance rate of dry ice vapors in the cage (not shown). The feedback-controlled temperature on the module was set to 33 °C. The temperature probe for the heating module was placed beneath the wire rack approximately centered over the cage.

**Figure 1:**
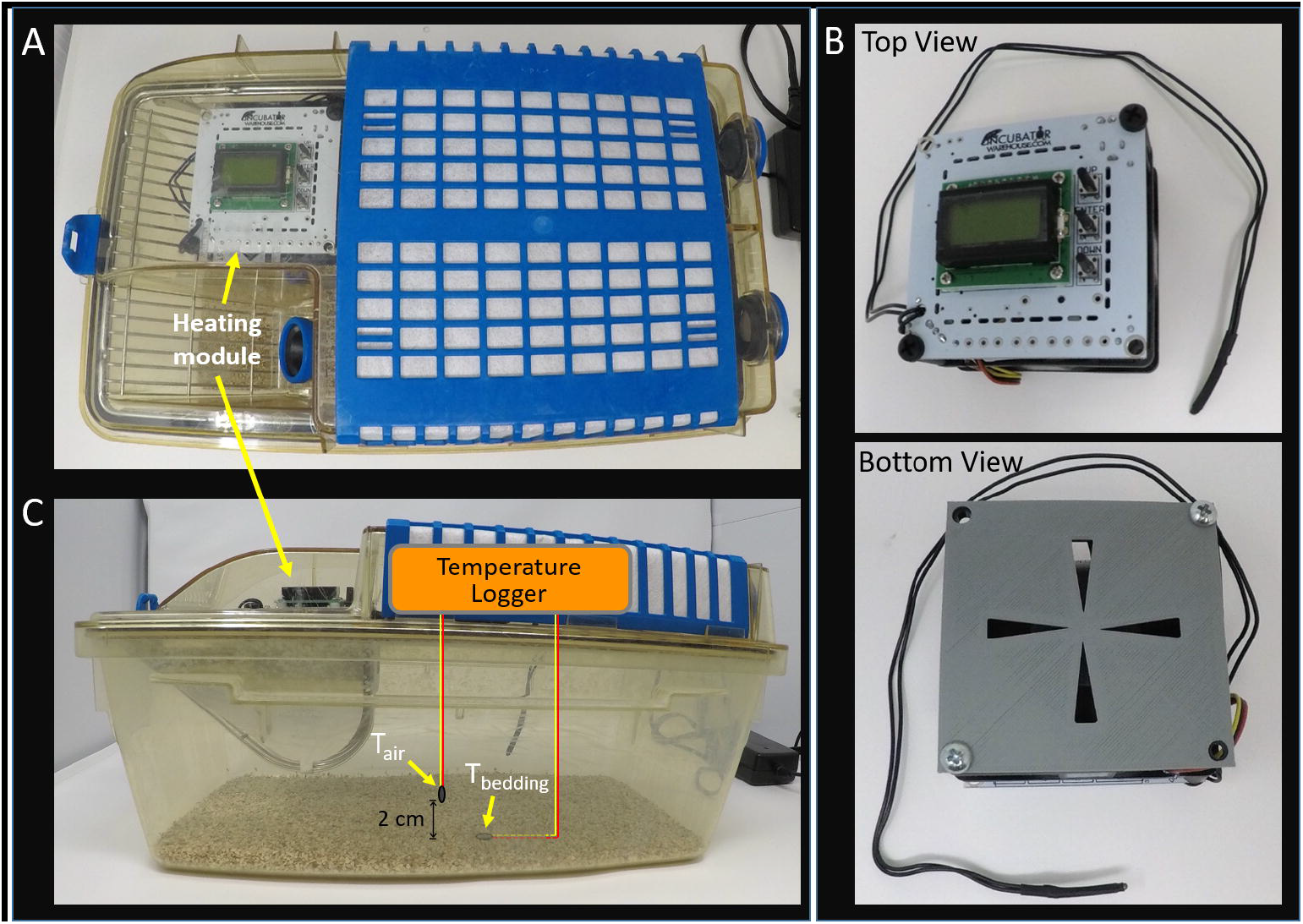
Assembly of warm ambient air cage (WAAC). (A) Top view of WAAC demonstrating location of heating module. (B) Heating module top view and bottom view with 3D-printed plastic fan baffle. (C) Side view of WAAC with indicated location of thermocouples for temperature characterization experiments. One thermocouple was placed 2 cm above the bedding (T_air_) and another placed under the top layer of the bedding (T_bedding_).

### Heating Pad Warmed Cage Design

A widely used method for providing post-surgical heat support to rodents is to place the cage half on and half off a heating pad (see **Fig 2A**). By doing so, it is assumed that the animal can warm on the heating pad side and then “escape” to the unheated side once the animal is either warm enough or too warm. To replicate this commonly used set up, we placed the cage half on a standard heating pad which was set to the “low” setting (Sun Beam).

**Figure 2:**
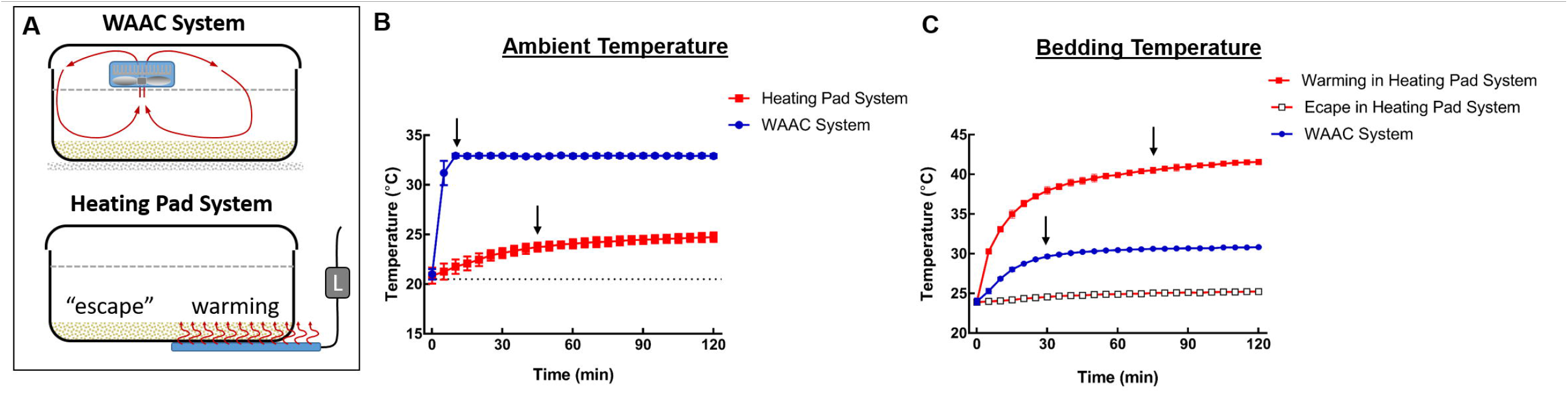
Comparison of heating characteristics for the WAAC and heating pad cage designs. (A) Drawing of the WAAC and heating pad systems. (B) Rate of ambient air warming (T_air_, measured above bedding). Time 0 represents the point where both systems were turned on. (C) Rate of bedding warming (T_bedding_, measured under top layer of bedding). For the heating pad system, bedding temperature was measured on the warming side (placed over the heating pad) and the unheated “escape” side (off the heating pad). The arrows represent the time for each system to reach within 1 °C of the steady-state temperature. Data are presented as mean ± SD from 3 separate measurements.

### Cage Temperature Characterization

For cage temperature characterization, we measured temperature at two points where a mouse would experience heat transfer – at the surface layer of the cage bedding and in the air just above the bedding. The location of these two thermocouples can be seen in **Fig 1D**. Temperature at these two locations was monitored by thermocouple and a temperature logger (HH806AU, Omega Engineering, Stamford, CT). Temperatures were recorded at 5 min intervals for up to two hours after turning on heating module or heating pad.

### Animal preparation and experimental groups

The experimental protocols were approved by the Institutional Animal Welfare Committee at the McGovern Medical School at UTHealth. A total of thirty-six male C57BL6/J mice (12 weeks old, Jackson Laboratory) were included in the study. The animals were allowed free access to food and water. Animals were randomly assigned to four experimental groups (n=9 mice, each) according to heat support method (WAAC or heating pad design) and severity of stroke model (mild or severe stroke). Mice were implanted with a small wireless capsule in the peritoneal cavity (Anipill, BodyCAP) (see **Supplemental Fig 1**). Upon activation, the capsule (∼16×8 mm) logged core temperature at 15 min interval. Mice were allowed at least 7 days to recover before additional experiments. Mice were also implanted with a small temperature transponder (IPTT-300, BioMedic Data Systems, Seaford, DE USA), which permitted the wireless measurement of body temperature. The transponder temperature reading could be performed through the wall of the cage and thus avoided the effect of stress (handling and restraint, rectal probe insertion) on body temperature. The wireless temperature measurement also eliminated the effect of repeated opening of the cage and exposure of the animal to cooler ambient air. Temperature measurements were made by an investigator blinded to the stroke severity grouping.

### Middle Cerebral Artery Occlusion (MCAo) Model of Mild and Severe Stroke

Mice were initially anesthetized by inhalation of 5% isoflurane in 30% O_2_/70% N_2_O, and then maintained with 1.5% isoflurane during the procedure. Rectal temperature was monitored and maintained at 37 °C by a feedback-controlled heating pad throughout the procedure (Omega Engineering). Core temperature (Tcore) was recorded by the Anipill capsule implanted in the peritoneal cavity and by temperature transponder (Implantable Programmable Temperature Transponder; IPTT-300, BioMedic Data Systems, Seaford, DE USA) which was implanted subcutaneously on the lower dorsolateral region of the mouse. This location placed the transponder in direct contact with the thin tissue layer surrounding the peritoneal cavity and thus provides good correlation with core body temperature(Feketa et al., 2014). The correlation between Anipill (intraperitoneal) and IPTT-300 (subcutaneous) temperatures was good (**Supplemental Fig 2**). Linear regression of the two temperature methods showed a nearly linear relationship, with a slope of 1.103 (P<0.0001).

Transient ischemic stroke was induced by occluding the middle cerebral artery (MCA) with a silicone-coated monofilament (MCAo model)(Fasipe Titilope et al., 2018). In brief, the occluder was introduced into the right external carotid artery (ECA), advanced back to the common carotid artery (CCA), and then advanced up the internal carotid artery (ICA) into the circle of Willis where it occluded the origin of the right MCA. The monofilament was left in place for 60 min and then removed to initiate reperfusion. Two occluder filaments were used to achieve the **mild** (0.23 mm diam, 2-3 mm coating length, 602323PK10Re, Doccol) or **severe** stroke injury (0.23 mm diam, 5-6 mm coating length, 602356PK10Re, Doccol). Once the occluder was in place, the mice were returned to the heat support recovery cage according to their assigned experimental group. Successful occluder placement was confirmed by neurologic deficit test (see below) during the stroke period. At 60 min, mice were again anesthetized and the monofilament withdrawn. Mice were again returned to their respective heat support recovery cages. Mice were evaluated for 3 days, at which point they were sacrificed for brain collection and analysis.

### Heat support during intra-ischemic and post-ischemic periods

Temperature support during the intra-ischemic period (time between monofilament placement and removal) and in the post-stroke period was provided by the WAAC or the heating pad design (described above). Mice were maintained in the heat support cages for 60 minutes during the occlusion period and for 3 hours following removal of the monofilament (i.e. reperfusion phase). Mice assigned to the heating pad system were placed in a region of the cage that was centered over the heating pad. Body temperature was measured via the wireless transponder and Anipill at 15 min intervals in the intra-ischemic period and at 30 min intervals in the post-ischemic period. After 3 hours of heat support, mice were transferred to standard cages (singly housed, no heat support). The housing facility ambient temperature was maintained at 25-26°C.

### Behavioral test (Neurologic deficit test)

Behavioral testing was performed to evaluate the degree of neurologic deficits based on a 12-point scoring system (12, maximal deficits; 0, no deficit)(Belayev et al., 1999). The test assessed postural reflex, forelimb placement by visual and tactile stimuli, and proprioceptive responses. We performed a neurologic deficits test just before reperfusion in order to confirm the successful occlusion. *A priori* exclusion criteria were set for mice *a)* not demonstrating a neurologic deficit score (NDS) of 12 at this time point, *b)* demonstrating evidence of vessel puncture by brain inspection or *c*) mouse barrel-rolling during MCAo. Two mice (one mouse in the heating pad system group with mild injury and one mouse in the WAAC system group with severe stroke) were excluded based on these criteria. Behavioral tests were additionally conducted at 24, 48 and 72 h after reperfusion.

### Histopathology

Mice were euthanized by overdose of isoflurane on post-stroke day 3 (PSD3). Brains were removed, sectioned at 1 mm thick coronal sections using a mouse brain matrix, and then incubated in 1% TTC (2,3,5-Triphenyltetrazolium Chloride, Sigma, in PBS) to evaluate infarct size. The stained sections were then fixed in 10% neutral buffered formalin before imaging with a stereomicroscope at 0.8X magnification (Olympus SZ-40). Cortical and subcortical infarcts in each section were outlined and measured by Image J. Infarct volume in the cortex and subcortex were calculated by multiplying infarct area in cortex or subcortex by sectional thickness and corrected for brain swelling.

Severity of hemorrhagic transformation (HT) was graded on a 0 to 5 point scale by an investigator blinded to grouping (see **Supplemental Fig 3**). The severity of HT was scored from 0 (none) to 5 (severe) for each section and then expressed as a cumulative total HT score.

### Statistical analysis

Data was expressed as a mean ± standard deviation (SD) or as median with interquartile range. The mean body temperature during intra-ischemic and post-ischemic periods was were analyzed by generalized estimating equation (GEE) method to address the within-subject correlation, followed by post-hoc group comparison at each time point. The homogeneity of variance in the two groups at each time point were evaluated by Bartlett test or Levene test depending on the normality. All p values were adjusted for multiple tests. NDS and HT score were analyzed by GEE method similarly, followed by post-hoc group comparison at each time point and comparisons between day 1, 2, 3 and baseline. All tests are adjusted for multiple tests. For single time measurements, groups were compared by two sample t test or Wilcoxon rank sum test. Data analyses were performed in GraphPad Prism 8 (San Diego, CA) and SAS 9.4 software (Cary, NC).

## Results

In total, 41 mice were used in this study. Eight mice died prior to the planned end point on PSD3 (72 hours) – 4 mice from the WAAC group and 4 mice from the heating pad groups **(see Table 1)**. The mild stroke injury resulted in very low mortality, with only one death in the heating pad and WAAC group prior to PSD3. The severe stroke injury group resulted in 3 deaths in the heating pad group (all on PSD1) and 4 deaths in the WAAC group (spread among PSD1-3). As noted in Methods, two mice (n=1, heating pad group with mild stroke; n=1, WAAC group with severe stroke) were excluded from the study. No differences were observed in initial weight or weight loss after stroke among heating pad and heating module cage groups (Supplemental Fig 4).

**Table 1:** Number of deaths before experiment end on post-stroke day 3 (PSD3). Number of deaths per group prior to PSD1-3. Two mice (n=1, heating pad group with mild stroke; n=1, WAAC group with severe stroke) were excluded due to surgical causes.

|  | Number of premature deaths |  |  |
| --- | --- | --- | --- |
| Group (n) | PSD1 | PSD2 | PSD3 |
| Heating pad, mild stroke (9) | 0 | 0 | 1 |
| Heating pad, severe stroke (12) | 3 | 0 | 0 |
| WAAC system, mild stroke (9) | 0 | 0 | 0 |
| WAAC system, severe stroke (11) | 1 | 1 | 2 |

### The WAAC system provides more rapid and precise cage temperature equilibration compared with the heating pad system

The rate of temperature stabilization was compared between the WAAC and heating pad cage systems (**Fig 2A**). To record temperature of bedding (T_bedding_) and ambient air (T_air_) from where a mouse would normally rest, one thermocouple was placed just under the top layer of bedding and another elevated two centimeters above the bedding. For the heating pad system, another thermocouple was placed in the bedding on the unheated “escape” side of the cage. Measurements were initiated at the same time that the respective heat support systems were powered on (time 0). The WAAC system heating module was set to 33 °C and the heating pad was adjusted to the “low” setting. The T_air_ in the WAAC reached the set point in under 10 min and remained extremely stable for the 2 hour period (**Fig 2B**). In contrast, the T_air_ in the heating pad cage responded more slowly, requiring 45 min to reach ±1 C of the steady-state temperature. The T_bedding_ increased to 30.85 °C in the WAAC, reaching ±1 °C of the equilibrated temperature in 30 min (**Fig 2C**). The T_bedding_ in the heating pad cage reached a higher temperature (41.5 ° C) and took 75 min to reach ±1 °C of the steady-state temperature. The bedding temperature on the “escape” side of the cage remained largely unchanged.

### The WAAC system provides more precise and stable control of mouse body temperature during the intra-ischemic and post-ischemic periods

Given the strong correlation between brain temperature and ischemic injury, it is critical to take steps to reduce the variability of this physiologic variable in experimental stroke studies(1999; Dirnagl, 2006; Fisher et al., 2009; Kahle and Bix, 2012; Percie du Sert et al., 2017). In this study, we measured body temperature as an easily accessible alternative to direct measurement of brain temperature in freely moving conscious mice. Brain temperature is influenced by multiple local factors (such as brain metabolic activity and blood flow), it can therefore differ in absolute temperature compared with core temperature or temporalis muscle temperature(DeBow and Colbourne, 2003; Jackson-Friedman et al., 1997; Kiyatkin et al., 2002). However, in non-stimulated conscious rodents, brain temperature generally parallels core temperature(DeBow and Colbourne, 2003; Kiyatkin, 2019, 2010), thus allowing body temperature measurements to be used as a reasonable approximation of brain temperature.

We measured Tcore in mice during and after 60 min of MCAO of either “mild” or “severe” stroke. The severity of stroke depended on the type of occluder used – the mild occluder had a 2-3 mm length of silicone tip, whereas the severe occluder had a 5-6 mm length. The longer silicone tip produces additional occlusion of the posterior cerebral artery (PCA) (**Supplemental Fig 5**), which results in the additional injury within the PCA territory. Heat support during surgery was provided by feed-back controlled heating pad and then by either WAAC or heating pad during the 60 minute intra-ischemic period and the first 3 hours of the post-ischemic (reperfusion) period.

In the mild stroke group, mean Tcore over the entire intra-ischemic period (15 to 60 min) was 36.4 ± 0.5 versus 35.9 ± 0.7 °C in WAAC system and heating pad groups, respectively. Mean differences were not significant (t test, P=0.119), nor was variance (Fisher’s F test, P=0.296). During the post-ischemic period, repeated measures analysis did not show differences by heating system at any time point. Furthermore, the variation at each time point was similar between groups. The mean Tcore during the reperfusion period (15 to 165 min) was 35.6 ± 0.4 versus 35.2 ± 0.4 °C, respectively (t test, P=0.131). As above, only mice surviving through PSD3 were included in the analysis. Mortality was low in the mild stroke groups (0 vs. 1, respectively) (see **Table 1**).

### The WAAC system reduces variability in functional outcome in severe stroke

Mice were evaluated by neurologic deficit score (NDS) at the end of the occlusion period (60 min) and at PSD1, 2 and 3. All mice demonstrated an NDS of 12 (full deficit) at the point just prior to removal of the occluder (**Fig 4**). Mice of all groups showed progressive functional improvement (lower NDS) through PSD3. The severe stroke group showed a correspondingly worse NDS compared with the mild stroke group. There was no overall group effect (across all days) by heat support system. Only mice that survived through PSD3 were included in the analysis.

In the severe stroke group, the WAAC system provided better and more consistent heat support during the intra-ischemic period (**Fig 3A**, top). The mean Tcore (from 15 to 60 min) for the WAAC system mice was 36.9 ± 0.3 °C versus 36.1 ± 1.3 °C in the heating pad system. While the mean temperatures were not statistically different (P=0.154, t test), the variance in the mean temperature was significantly less in the WAAC group (Bartlett’s test, p =0.004). During the post-ischemic phase, the WAAC system produced significantly better Tcore maintenance (and closer to 37 °C) and lower Tcore variability compared with the heated cage system for the severe stroke group mice (**Fig 3B**, top). Repeated measures analysis showed significantly improved temperature maintenance during the post-ischemic period, with statistical significance at 15, 30, 45, 75, 105 135, and 165 minutes after reperfusion (P<0.05, Bartlett’s test). The mean Tcore over the entire reperfusion period was 35.9 ± 0.4 °C versus 34.2 ± 1.7 °C for WAAC and heating pad groups, respectively. The mean Tcore differed significantly (p=0.029, t-test), and the variance was significantly lowered in the WAAC group (p=0.004, Barlett’s test). To avoid skewing the data, only mice that survived through PSD3 were included in the temperature analysis. Mortality rates through PSD3 were similar between groups (4 vs. 3, respectively). However, while mortality occurred exclusively in the first 24 hours in the heating pad group, mortality was spread across the 72 hour period in the WAAC group (see **Table 1**).

**Figure 3:**
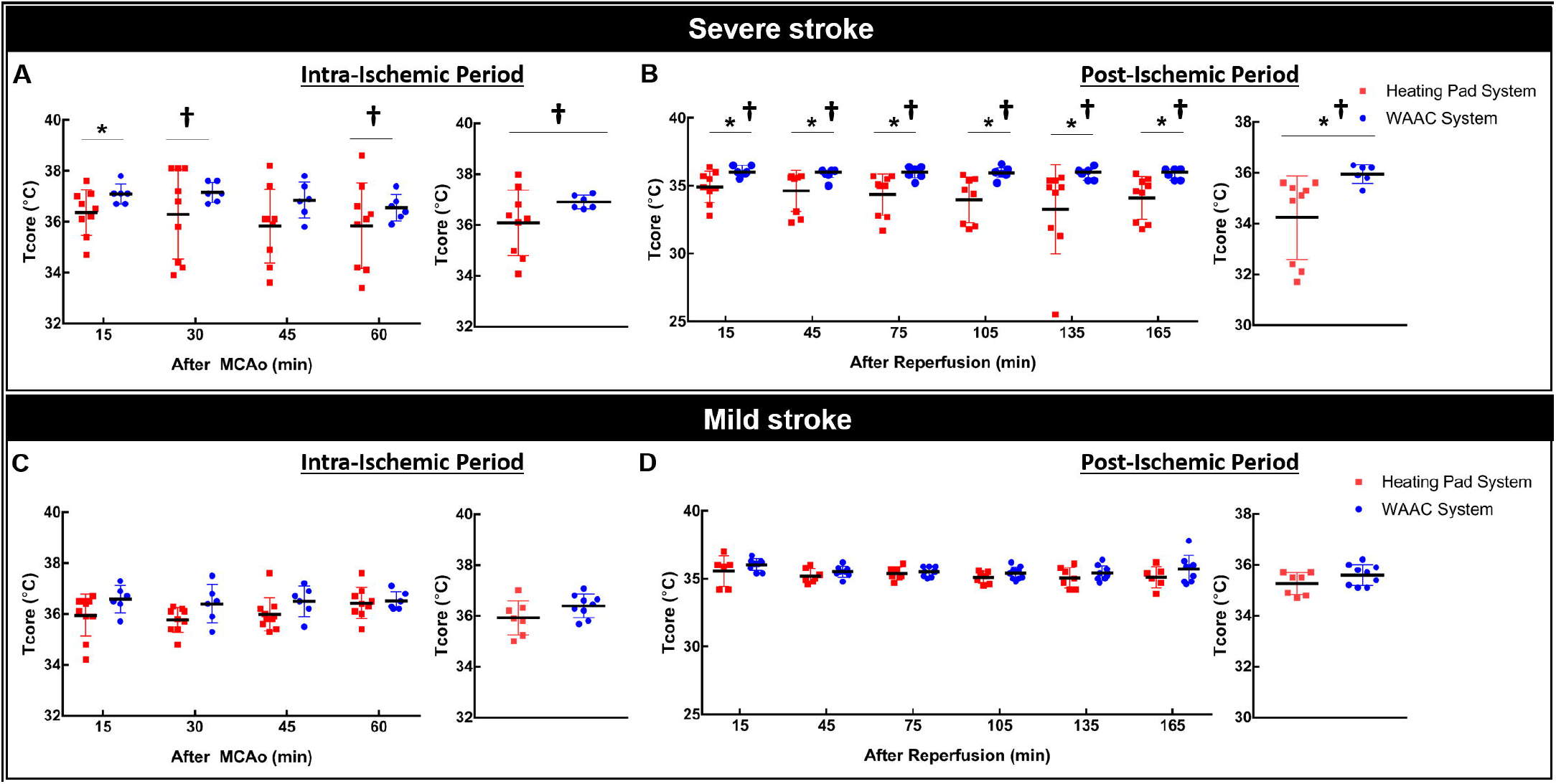
Comparison of core temperature (Tcore) management by heat support system during the intra-ischemic and post-ischemic phase for mice that underwent the severe stroke protocol (A,B) or the mild stroke protocol (C,D). (A,C) Summary of intra-ischemic Tcore for WAAC and heating pad groups at different time points during the occlusion (left) and the mean Tcore over the entire occlusion period (right). The † indicates significantly lower variance in the WAAC group (GEE method). (B,D) Summary of post-ischemic Tcore for the WAAC and heating pad groups at different time points (left) and over the entire reperfusion period (right). The * indicates group differences in mean temperature. All-time points in the WAAC system with severe stroke reached statistical significance. The † indicates significantly lower variance in the WAAC group (Bartlett test or Levene’s test, P values were adjusted for multiple tests). All data are presented as mean ± SD.

**Figure 4:**
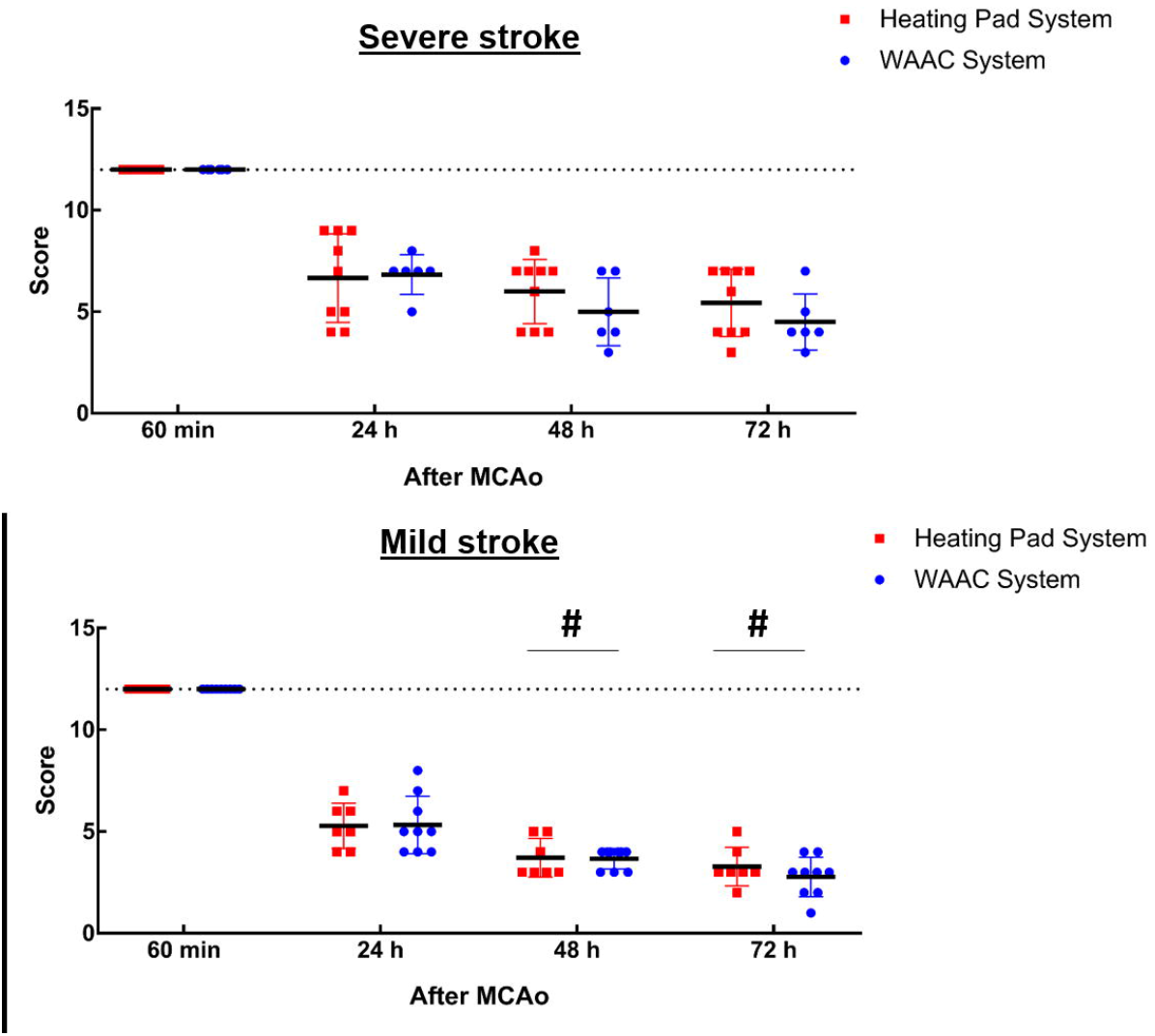
Comparison of neurological deficit score (NDS) by heat support system through PSD3. Data are presented as mean ± SD; the # reflects significant intra-group difference from PSD1 (GEE method).

**Figure 5:**
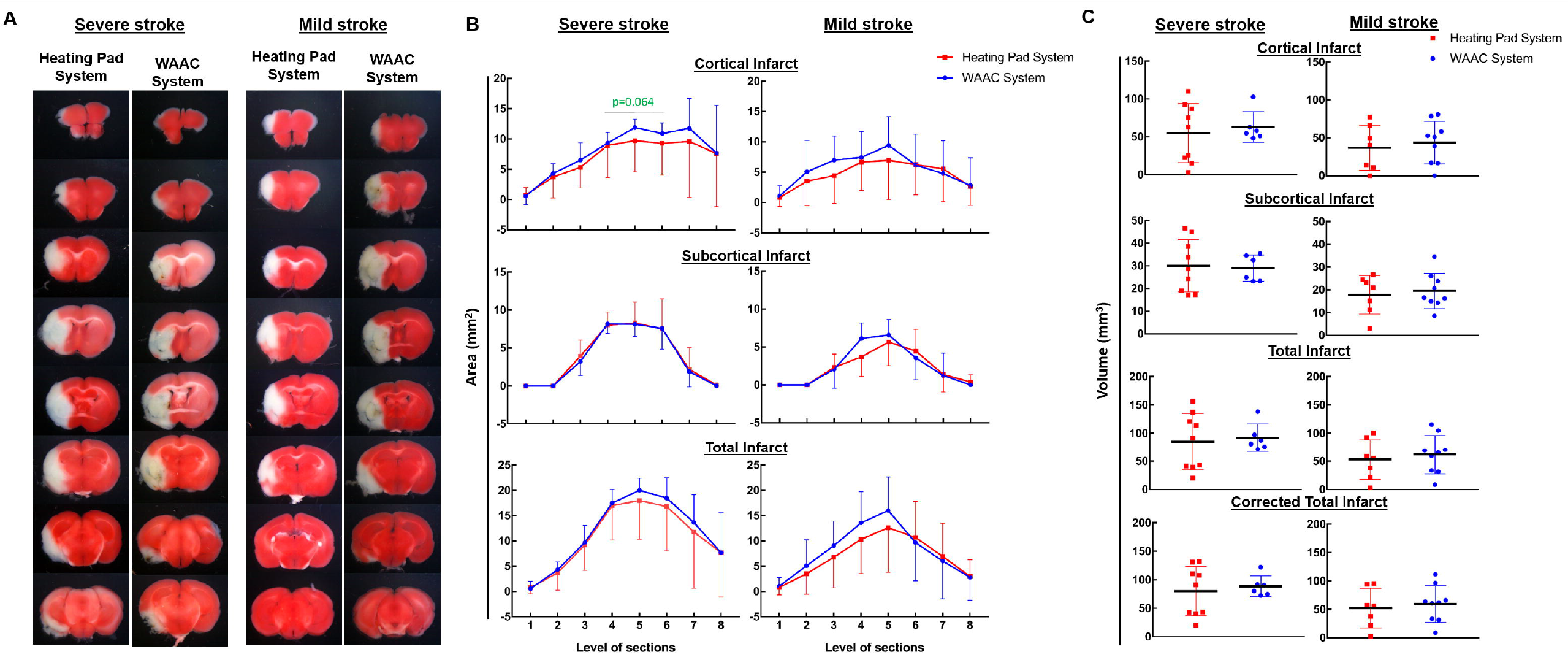
Comparison of infarct at PSD3 between heating systems. (A) Representative TTC stained brain sections for WAAC and heating pad systems for severe and mild stroke models. Severe stroke brains show infarct in posterior sections, reflecting injury to both the middle cerebral artery (MCA) and posterior cerebral artery (PCA) territories. Mild stroke brains show injury to the MCA territory only. (B) Summary of infarct area by brain section between groups. Areas are reported as mean ± SD for cortical infarct, subcortical infarct, and combined total infarct. (C) Summary of infarct volume between groups, plotted as all points, mean volume, and SD for cortical, subcortical, total, and total corrected for swelling infarct volume. No group differences were found; however, SD values were consistently lower in the WAAC heat support group with severe stroke.

### Effect of WAAC system on infarct volume variability

Brains were harvested at PSD3 and analyzed for infarct volume as cortical infarct, subcortical infarct, and total infarct (with and without correction for swelling) (**Fig 5**). In the severe stroke group, no differences were found in mean infarct volumes between heat support systems. However, the variances in cortical infarct areas at level 4,5 and 6 approached significance, with P=0.064, 0.064, and 0.064 respectively (**Fig 3B**, top). Infarct volumes in cortical, subcortical, total and corrected infarct volume (in mm^3^ for WAAC and heating pad, respectively) were: 63.0 ± 20.2 vs. 54.9 ± 39.0 (cortex), 29.0 ± 5.8 and 30.1 ± 11.5 (subcortex), 92.0 ± 24.4. vs. 58.0 ± 50.0 (total), and 88.7 ± 18.2 vs. 80.1 ± 43.3 (corrected total). The variability of the infarct volume data (SD values) was consistently lower in the WAAC group (1.9 to 2.4 fold reduced), but did not reach statistical significance in these limited group sizes (Levene’s test; P=0.1880, 0.0932, 0.1410, and 0.1030, respectively). In the mild stroke group, the infarct was predominantly evident in the more anterior brain sections and was reduced compared with the severe stroke group. Within the mild stroke group, neither the infarct volumes nor the infarct volume variability differed by heat support system. Infarct volumes in cortical, subcortical, total and corrected infarct volume (in mm^3^ for WAAC and heating pad, respectively) were: 43.8 ± 28.1 vs. 36.8 ± 11.2 (cortex), 19.5 ± 7.7 vs. 17.9 ± 8.5 (subcortex), 62.0 ± 34.3 vs. 53.1 ± 35.5 (total), and 59.5 ± 32.0 vs. 52.4 ± 34.7 (corrected total).

### Effect of WAAC system on hemorrhagic transformation score

Hemorrhagic transformation (HT) was evaluated at PSD3 by gross inspection of high-resolution images of the TTC-stained sections. Representative sections showing the presence of hemorrhage (yellow arrows) in severe and mild stroke brains are shown in **Fig 6A**. HT was scored (0-5 scale) by blinded investigator for each section and then summed to obtain a total score for each brain. There was no overall group effect of heat support system in either severe and mild stroke models (panels B and C), however, there were individual differences in the severe stroke group at sections 2 and 7 and in the mild stroke group at section 5.

**Figure 6:**
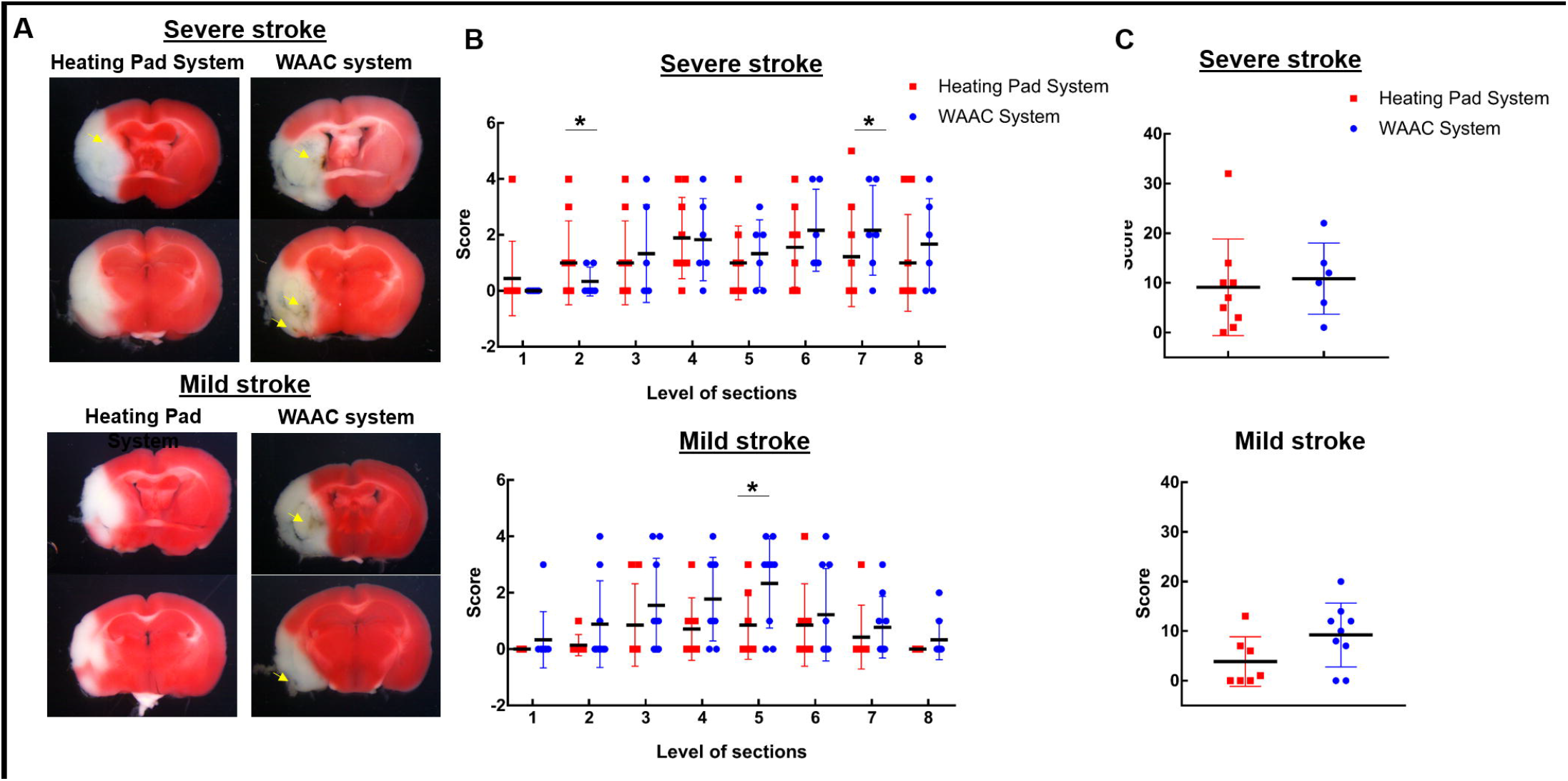
Comparison of hemorrhagic transformation (HT) at PSD3 between heating systems. (A) Representative TTC stained brains showing evidence of hemorrhage in the infarct region (yellow arrows). (B) Summary of HT score (0-5) by brain section for severe stroke (top) and mild stroke (bottom) for both heat support systems. HT score is presented as mean ± SD. The HT score was significantly increased in the WAAC system mice of the severe (at sections 2 and 7) and mild (at section 5) stroke groups (post-hoc group comparison by GEE method, adjusted for multiple testing). (C) Summary of combined HT score, with data presented as in B. No overall group differences were evident.

## Discussion

The primary findings of this study are *1*) the warmed ambient air cage (WAAC) system provides more precise control of post-stroke mouse body temperature compared with traditional heating pad warming system and *2*) more precise control of post-stroke core temperature reduces variability in some experimental measures in more severely injured mice. These findings re-affirm the importance of tightly controlling body temperature during the intra-ischemic and post-ischemic periods to reduce variability in experimental stroke outcomes. These studies additionally describe the fabrication of a low-cost cage warming system that can be assembled and applied in any stroke laboratory.

Our findings showed that the WAAC system was significantly more stable than the heating pad system, reaching steady-state cage temperature more quickly and then maintaining cage temperature very precisely. Although our study did not require frequent opening of the lid, we found that the WAAC system was able to quickly recover temperature after lid opening within (2-3) minutes (**Supplemental Fig 6**). The WAAC system is easy to assemble and the components are relatively inexpensive. With the cost of the heater/circulator unit at <$60 USD, a laboratory can assemble multiple recovery cages to accommodate high-throughput studies. Although not shown, the heater system also provided good temperature control for a rat cage.

We compared our WAAC system to a widely used form of post-surgical warming (cage on heating pad) to evaluate its effectiveness on animal temperature stability as well as variability in stroke outcomes. We used a common model of stroke (MCAo) with two versions of commercially available occluder devices. The occluder devices consisted of a 6-0 monofilament with the occluding tip coated with silicone to a diameter of 0.21 mm. In the C57BL/6 strain of mice, the coating length affects whether the occluder blocks flow to the middle cerebral artery (MCA) alone (i.e. 2-3 mm occluder) or to the MCA and the posterior cerebral artery (PCA) (i.e. 5-6 mm occluder) (see **Supplemental Fig 5**). As a result, these occluders produce either a more mild stroke (2-3 mm, MCA territory only) or a more severe stroke (5-6 mm, MCA and PCA territory).

Mice and other mammals maintain body temperature through a balance of metabolic heat production and temperature exchange with the environment. During times of reduced metabolic activity (e.g. anesthesia, stroke, etc.), this balance is shifted and body temperature can fall. This drop in temperature can be both rapid and extreme in mice due to their lower mass to body surface area relationship(Gordon, 2012). In relation to body warming, two important temperature ranges are the thermoneutral zone (TNZ) and the ambient temperature range for normothermia. The TNZ reflects the ambient temperature range over which mice are metabolically neutral (i.e. increased metabolic activity is not needed to maintain core temperature). In mice, the TNZ varies according to conditions, but is typically 28-32 °C(Gordon, 2012). The ambient temperature range for normothermia reflects the point at which further increased temperature results in an increase in core temperature(Gordon, 2012). The ambient temperature of the WAAC system was set to 33 °C, which falls just above the thermoneutral zone (TNZ) in mice of ∼28-32 °C and within the ambient temperature range for normothermia (32 – 34 °C). By providing an elevated ambient temperature, our WAAC system compensates for the loss in metabolic heat production by the post-surgery/post-stroke mice, and thus allows the mice to maintain a more consistent and normothermic body temperature. Furthermore, by setting the ambient temperature (33 °C) only slightly above the TNZ, mouse temperature stays within the normothermia range – and importantly, below the temperature that results in *hyper*thermia.

Our choice to measure body temperature (instead of direct brain temperature) was based on practical issues, and thus on what could be easily applied in stroke studies in any laboratory. While measurement and control of brain temperature would be ideal, its direct measurement requires insertion of a temperature probe into the brain. This procedure adds cost, an additional surgical procedure, and produces brain tissue damage. Given that intra-ischemic and post-ischemic *body* temperature is not currently widely reported in experimental stroke studies, the expectation that the field would adopt routine *brain* temperature measurements is low. While not a perfect reflection of absolute brain temperature, core temperature has been shown to track reasonably well with brain temperature in non-stimulated conditions(DeBow and Colbourne, 2003; Jackson-Friedman et al., 1997; Kiyatkin and Wise, 2002). In mice, the small subcutaneous transponders show good correlation with intraperitoneal temperature(Hankenson et al., 2018; Kort et al., 1998) (**Supplemental Fig 2**), and thus offer a simple and inexpensive option for wirelessly monitoring core temperature. The transponders used in the present study cost approximately $10 USD per unit and can be sterilized and reused several times. The wand reader is the most expensive item, costing around $2500 USD. However, since brain temperature is one of the most influential variables in brain injury, tightening this variable through more precise heat support should enable stroke studies to be performed with fewer mice while maintaining equal or better statistical power. As an example, using infarct volume as the measured variable in a power analysis calculation, the WAAC system would reduce the number of mice needed by >4 fold compared with the heating pad system (based on a 50% change in means).

Our findings showed that the primary benefit of the WAAC system was in the mice which underwent the severe stroke protocol. By comparison, the traditional heating pad system mice in the severe stroke group demonstrated larger deviation of body temperature from normothermia compared with the WAAC mice. In addition, the heating pad mice showed greater variation in temperature, evaluated in both the intra-ischemic and post-ischemic periods. The mild stroke mice showed no statistically significant effect by heat support method. The reason for the difference between stroke injuries is presumably due to a greater reduction in activity-dependent heat production in the more severely injured animals. Note that the hypothalamus, a key brain region in thermoregulation(Morrison and Nakamura, 2011; Tan and Knight, 2018), was not injured in either stroke protocol (see **Fig 5**). From these findings, the WAAC system should also provide improved temperature control in other stroke models where stroke injury is more severe, such as in aged mice or mice with exacerbating co-morbidities.

## Limitations

As noted above, our studies utilized core temperature as a surrogate of brain temperature. While brain temperature often parallels body temperature in conscious rodents, absolute brain temperature can range from ∼1 °C above or below core temperature(DeBow and Colbourne, 2003; Kiyatkin et al., 2002) and can further deviate during specific physiological challenges (e.g. tail pinch, social stress) and pharmacological challenges (Kiyatkin and Wise, 2002). The brain itself is a significant source of metabolic heat, and thus brain temperature can further dissociate from core temperature at times when increased brain activity is not adequately accounted for by increased heat removal via the circulatory system. Measurement of temporalis muscle temperature is sometimes used as an approximation of brain temperature in rodents, particularly in rats(Jackson-Friedman et al., 1997; Kiyatkin, 2010). The temporalis muscle is physically associated with the head and is also supplied by the carotid artery. Thus, the temporalis muscle temperature is considered to be a reflection of both the temperature of arterial blood supplying the brain as well as the underlying brain, which is separated by the skull. Unfortunately, the MCAo stroke model involves permanent ligation of the external carotid artery, which is the branch of the carotid that supplies much of the scalp and the temporalis muscle(Ku and Choi, 2012; Vaas et al., 2017). In addition, the temporalis muscle in mice is of significantly less mass than in rats(Liu et al., 2009). These factors make the temporalis muscle less ideal (or at least far more complicated) as a proxy of brain temperature in mice. Our approach based on providing tighter core temperature control, while not without limitations, appears to have nevertheless produced the desired effect of reducing variability in stroke outcome.

In summary, we describe a low-cost cage warming system (WAAC) that reflects a significant improvement over the widely used “cage on a heating pad” approach. The WAAC T_air_ and T_bedding_ can be more precisely controlled, which leads to tighter body temperature control and reduced variability in stroke outcome. In addition, the WAAC system prevents accidental *hyper*thermia, as can occur with heating pads without feedback control systems. These refinements in intra-ischemic and post-ischemic temperature management are consistent with the goals of the Stroke Therapy Academic Industry Roundtable (STAIR), the Ischaemia Models: Procedural Refinements Of in Vivo Experiments (IMPROVE), and other recommendations for improving translation of stroke findings(1999; Dirnagl, 2006; Kahle and Bix, 2012; Percie du Sert et al., 2017). The described approach of wireless temperature monitoring and precise temperature support could easily be applied in stroke research laboratories.

## Acknowledgements

None.

## Funding acknowledgements

Funding for the project included AHA Postdoctoral Fellowship (19POST34380074) to SH and NIH NS094280, NS 096186 to SPM.

## Author contribution statement

Contribution consisted of conception of the project (SH, ML, SPM), conducting experimental studies (SH, JH, ML, JS), summary and analysis of data (SH, LZ, SPM), and writing the manuscript (SH, SPM).

## Supplemental Figures

**Supplemental Figure 1:** Implantable temperature sensors (Anipill and IPTT-300) and respective detectors for wireless measurement of mouse body temperature. (A) Anipill sensor programing unit. (B) Anipill wireless receiver unit. (C) Anipill implantable temperature sensor with wireless and data logging capabilities. (D) IPTT reader wand. (E) IPTT-300 hypodermic delivery unit. (F) IPTT-300 temperature transponder.

**Supplemental Figure 2:** Correlation between Anipill and IPTT-300 temperature measurements. The Anipill was placed in the peritoneal cavity and the IPTT-300 was placed sub-cutaneously in the same mouse. Temperature was measured at different time points during the post-ischemic period from 27 mice. The data is presented as the full range (left) and the 34-37 °C range (right). A linear regression with 95% confidence intervals is shown on both plots, generated from the full range of data.

**Supplemental Figure 3:** Hemorrhagic transformation (HT) scoring system. Representative images for a given score as described by the size of hemorrhage (small versus large) and the relative area of hemorrhage within the infarct. Scoring is done by an investigator blinded to group.

**Supplemental Figure 4:** Summary of physiologic variables. Day 0 reflects the pre-surgery values. No significant differences by group.

**Supplemental Figure 5:** Microfil infused brain of adult C57BL/6 mouse showing the approximate occlusion region (blue region) of the 2-3 mm occluder (A) and the 5-6 mm occluder (B). The silicone tip of the occluder is introduced into the circle of Willis via the internal carotid artery (ICA) and advanced past the ostium of the middle cerebral artery (MCA). The 2-3 mm occluder blocks flow to the MCA, whereas the longer 5-6 mm occluder additionally blocks flow to the posterior cerebral artery (PCA). Note that even when the origin of the PCA is blocked, some flow to the PCA may persist from the posterior communicating artery (PCom), which delivers flow from the vertebrals/basilar. In the C57BL/6, the PCom is typically small in diameter and can be unilateral or even absent in many mice. In the 2-3 mm occluder mice, flow to the PCA is maintained via the ICA since the common carotid artery is kept patent during the occlusion period.

**Supplemental Figure 6:** Demonstration of rapid ambient air temperature (Tair) recovery following cage lid removal in the WAAC system. After reaching steady-state temperature, the cage lid was removed for 10 seconds and then replaced. Tair is presented as mean ± SD from 4 separate measurements.

## References

Recommendations for Standards Regarding Preclinical Neuroprotective and Restorative Drug Development. Stroke, 1999; 30: 2752-8.

Belayev L, Busto R, Zhao W, Fernandez G, Ginsberg MD. Middle cerebral artery occlusion in the mouse by intraluminal suture coated with poly-l-lysine: neurological and histological validation. Brain Research, 1999; 833: 181-90.

Buchan A, Pulsinelli W. Hypothermia but not the N-methyl-D-aspartate antagonist, MK-801, attenuates neuronal damage in gerbils subjected to transient global ischemia. The Journal of Neuroscience, 1990; 10: 311-6.

Cao Z, Balasubramanian A, Marrelli SP. Pharmacologically induced hypothermia via TRPV1 channel agonism provides neuroprotection following ischemic stroke when initiated 90 min after reperfusion. Am J Physiol Regul Integr Comp Physiol, 2014; 306: R149–R56.

Cao Z, Balasubramanian A, Pedersen SE, Romero J, Pautler RG, Marrelli SP. TRPV1-mediated Pharmacological Hypothermia Promotes Improved Functional Recovery Following Ischemic Stroke. Scientific reports, 2017; 7: 17685-.

DeBow S, Colbourne F. Brain temperature measurement and regulation in awake and freely moving rodents. Methods, 2003; 30: 167-71.

Dirnagl U. Bench to Bedside: The Quest for Quality in Experimental Stroke Research. Journal of Cerebral Blood Flow & Metabolism, 2006; 26: 1465-78.

Fasipe Titilope A, Hong S-H, Da Q, Valladolid C, Lahey Matthew T, Richards Lisa M, Dunn Andrew K, Cruz Miguel A, Marrelli Sean P. Extracellular Vimentin/VWF (von Willebrand Factor) Interaction Contributes to VWF String Formation and Stroke Pathology. Stroke, 2018; 49: 2536-40.

Feketa VV, Zhang Y, Cao Z, Balasubramanian A, Flores CM, Player MR, Marrelli SP. Transient Receptor Potential Melastatin 8 Channel Inhibition Potentiates the Hypothermic Response to Transient Receptor Potential Vanilloid 1 Activation in the Conscious Mouse. Critical Care Medicine, 2014; 42: e355.e63.

Fisher M, Feuerstein G, Howells DW, Hurn PD, Kent TA, Savitz SI, Lo EH. Update of the Stroke Therapy Academic Industry Roundtable Preclinical Recommendations. Stroke, 2009; 40: 2244-50.

Gordon CJ. Thermal physiology of laboratory mice: Defining thermoneutrality. Journal of Thermal Biology, 2012; 37: 654-85.

Hankenson FC, Marx JO, Gordon CJ, David JM. Effects of Rodent Thermoregulation on Animal Models in the Research Environment. Comparative medicine, 2018; 68: 425-38.

Jackson-Friedman C, Lyden PD, Nunez S, Jin A, Zweifler R. High Dose Baclofen Is Neuroprotective but Also Causes Intracerebral Hemorrhage: A Quantal Bioassay Study Using the Intraluminal Suture Occlusion Method. Experimental Neurology, 1997; 147: 346-52.

Kahle MP, Bix GJ. Successfully Climbing the “STAIRs”: Surmounting Failed Translation of Experimental Ischemic Stroke Treatments. Stroke Res Treat, 2012; 2012: 374098-.

Kiyatkin EA. Brain temperature and its role in physiology and pathophysiology: Lessons from 20 years of thermorecording. Temperature, 2019; 6: 271-333.

Kiyatkin EA. Brain temperature homeostasis: physiological fluctuations and pathological shifts. Front Biosci (Landmark Ed), 2010; 15: 73-92.

Kiyatkin EA, Brown PL, Wise RA. Brain temperature fluctuation: a reflection of functional neural activation. European Journal of Neuroscience, 2002; 16: 164-8.

Kiyatkin EA, Wise RA. Brain and Body Hyperthermia Associated with Heroin Self-Administration in Rats. The Journal of Neuroscience, 2002; 22: 1072-80.

Kort WJ, Hekking-Weijma JM, TenKate MT, Sorm V, VanStrik R. A microchip implant system as a method to determine body temperature of terminally ill rats and mice. Lab Anim, 1998; 32: 260-9.

Ku T, Choi C. Noninvasive optical measurement of cerebral blood flow in mice using molecular dynamics analysis of indocyanine green. PLoS One, 2012; 7: e48383.e.

Liu L, Yenari MA. Therapeutic hypothermia: neuroprotective mechanisms. Frontiers in bioscience: a journal and virtual library, 2007; 12: 816-25.

Liu S, Zhen G, Meloni BP, Campbell K, Winn HR. RODENT STROKE MODEL GUIDELINES FOR PRECLINICAL STROKE TRIALS (1ST EDITION). J Exp Stroke Transl Med, 2009; 2: 2-27.

Morrison SF, Nakamura K. Central neural pathways for thermoregulation. Front Biosci (Landmark Ed), 2011; 16: 74-104.

Percie du Sert N, Alfieri A, Allan SM, Carswell HV, Deuchar GA, Farr TD, Flecknell P, Gallagher L, Gibson CL, Haley MJ, Macleod MR, McColl BW, McCabe C, Morancho A, Moon LD, O’Neill MJ, Pérez de Puig I, Planas A, Ragan CI, Rosell A, Roy LA, Ryder KO, Simats A, Sena ES, Sutherland BA, Tricklebank MD, Trueman RC, Whitfield L, Wong R, Macrae IM. The IMPROVE Guidelines (Ischaemia Models: Procedural Refinements Of in Vivo Experiments). Journal of Cerebral Blood Flow & Metabolism, 2017; 37: 3488-517.

Tan CL, Knight ZA. Regulation of Body Temperature by the Nervous System. Neuron, 2018; 98: 31-48.

Vaas M, Ni R, Rudin M, Kipar A, Klohs J. Extracerebral Tissue Damage in the Intraluminal Filament Mouse Model of Middle Cerebral Artery Occlusion. Front Neurol, 2017; 8: 85.

