## Supplementary figures and images for "A low-cost mouse cage warming system provides improved intra-ischemic and post-ischemic body temperature control – application for reducing variability in experimental stroke studies"

### Supplemental Figure 1

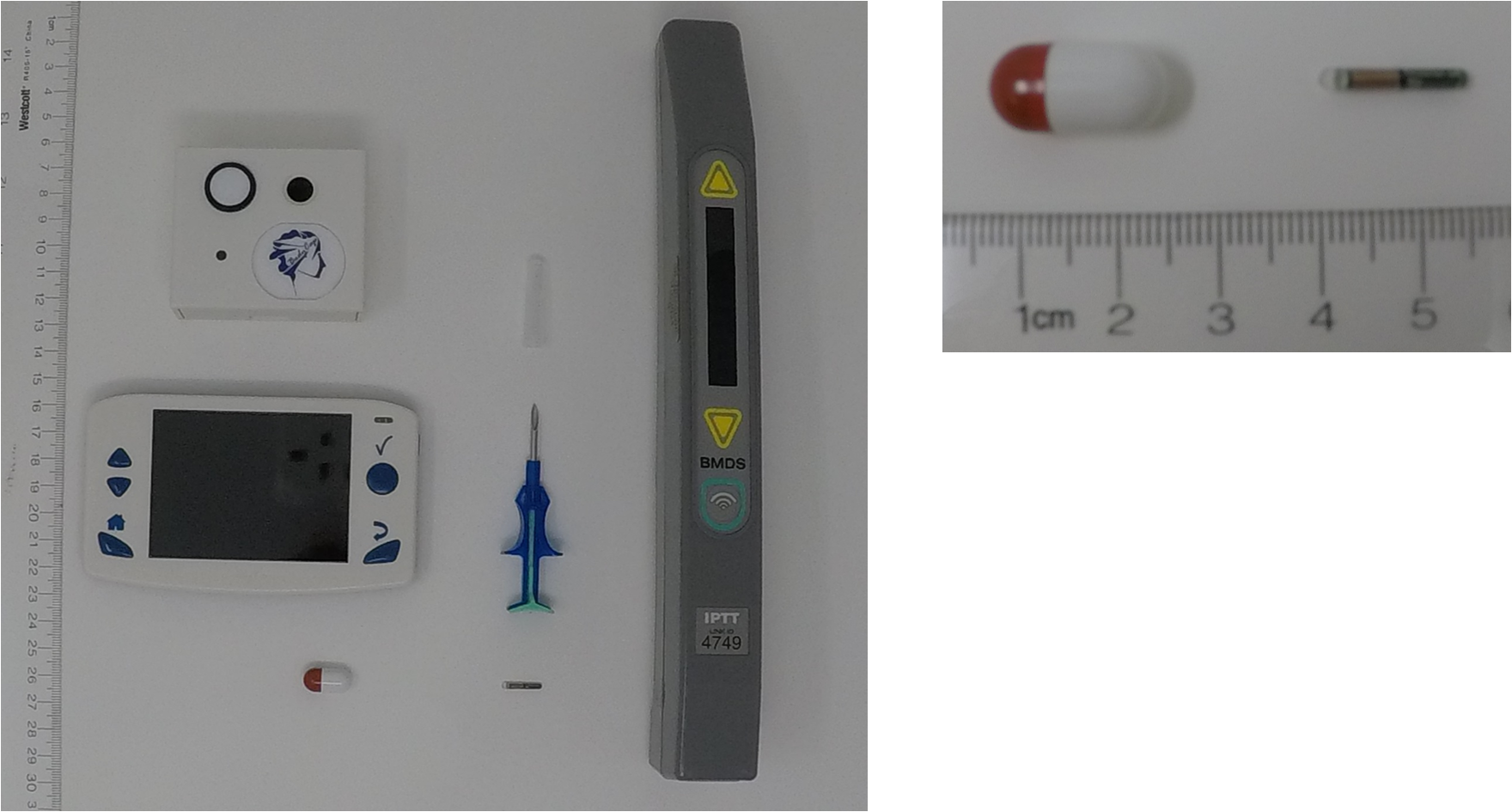

### Supplemental Figure 2

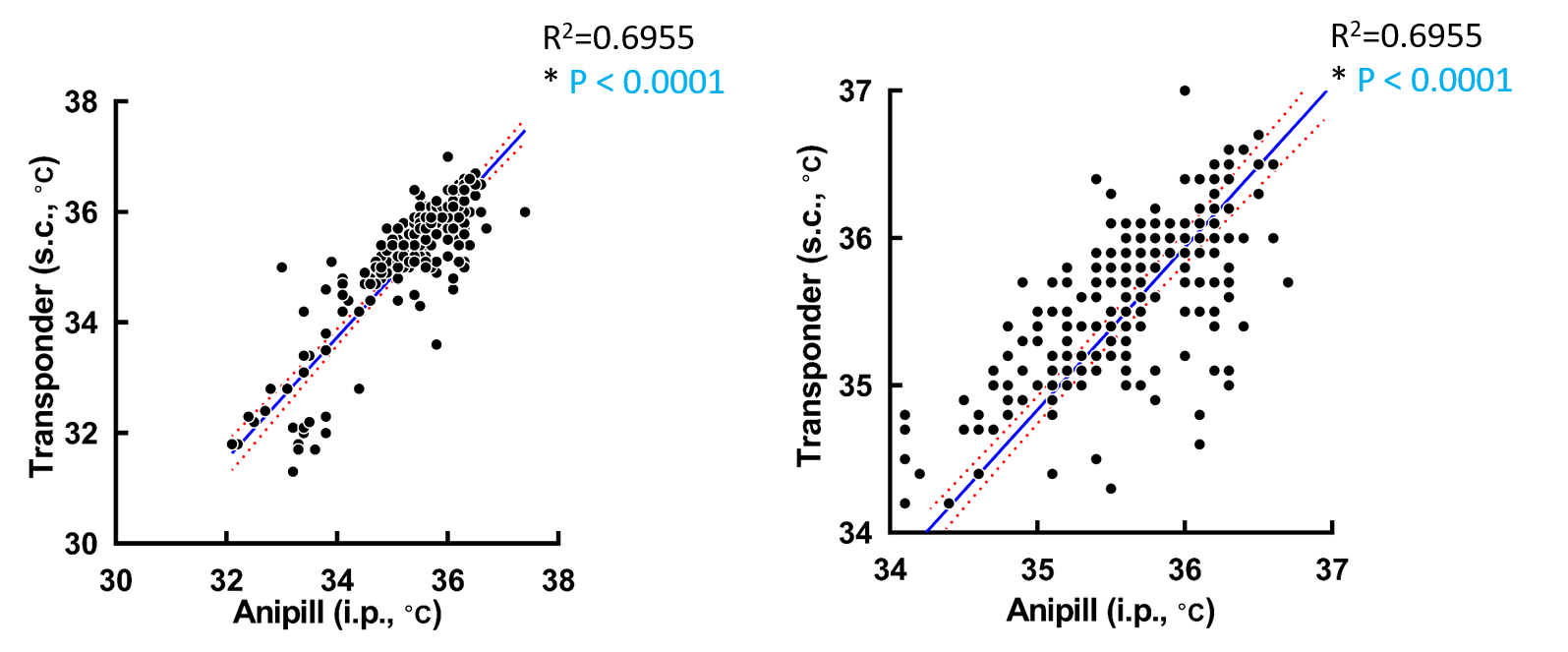

### Supplemental Figure 3

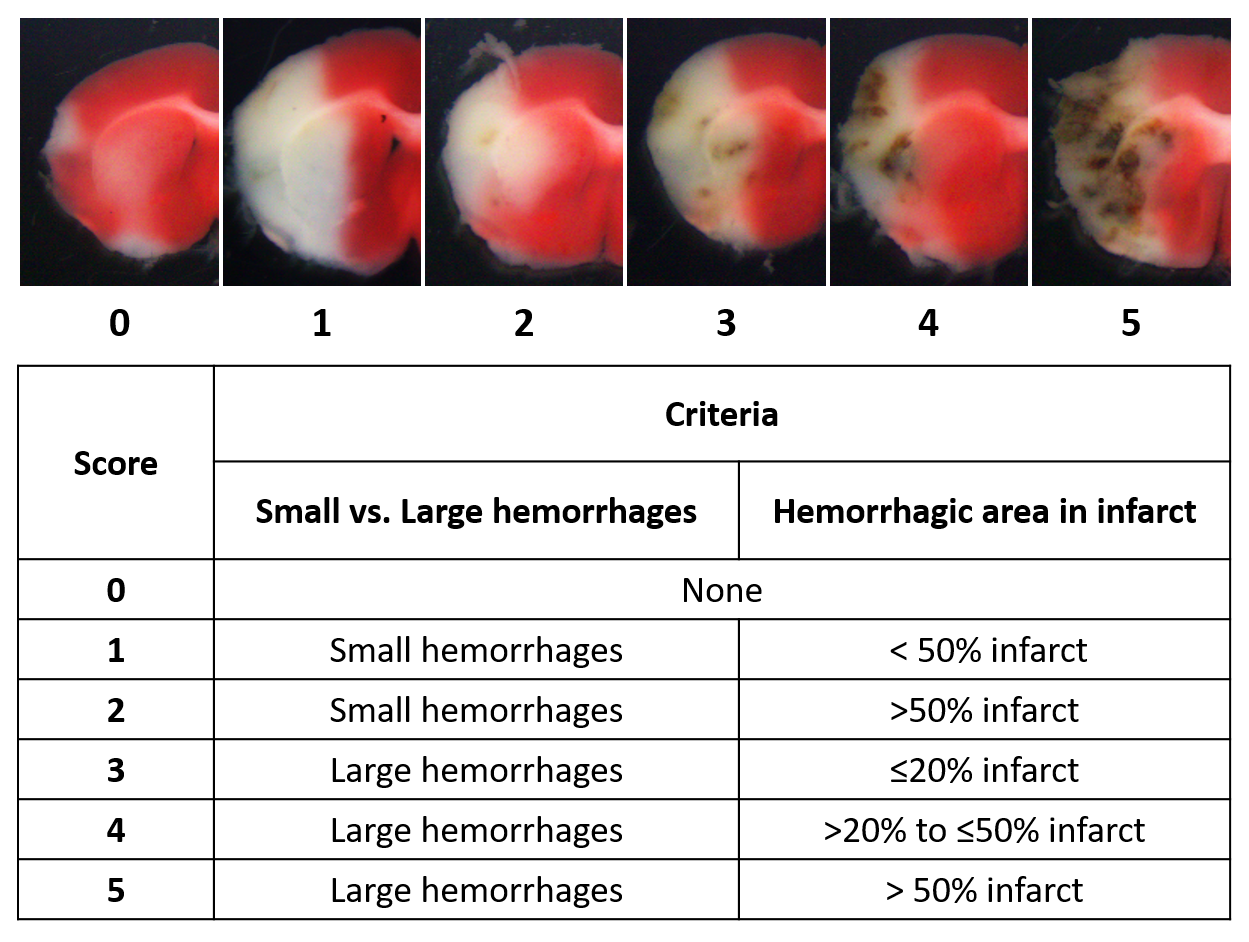

### Supplemental Figure 4

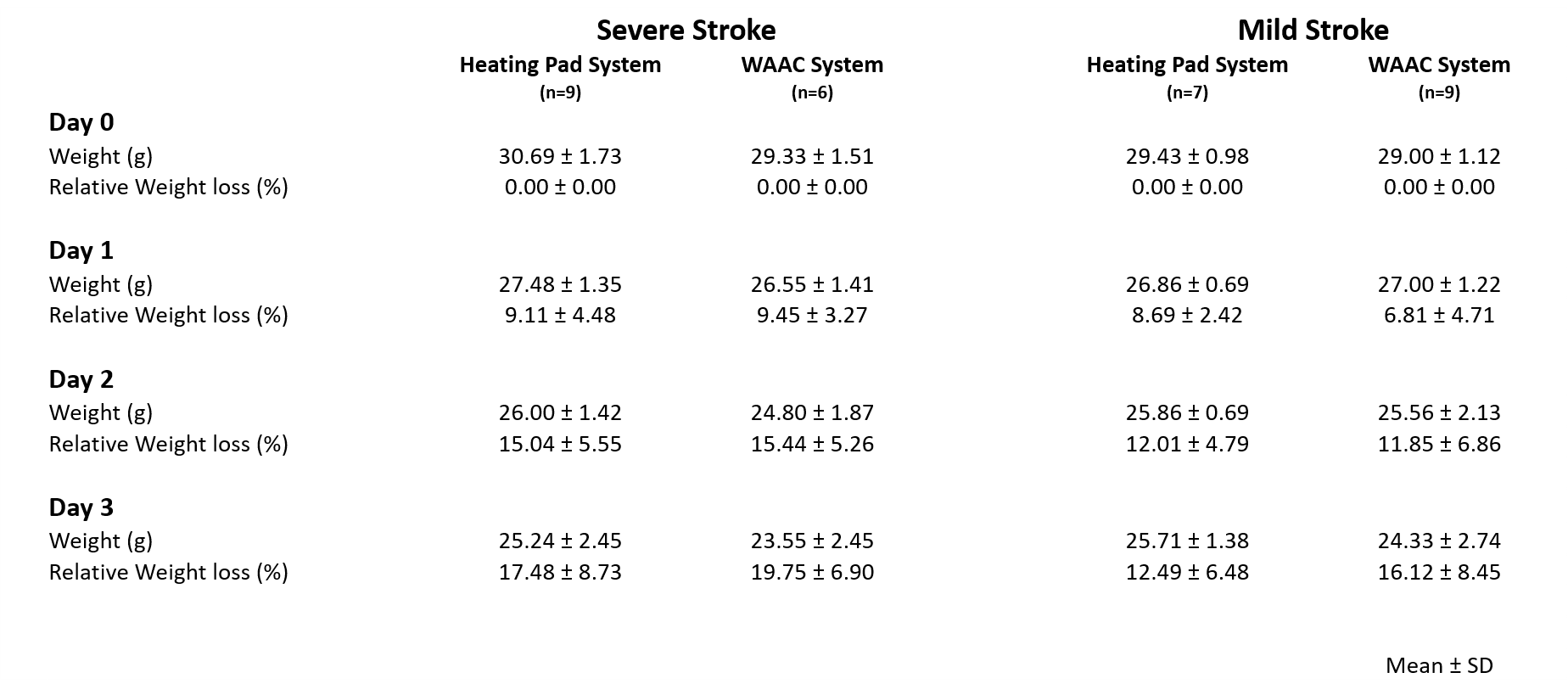

### Supplemental Figure 5

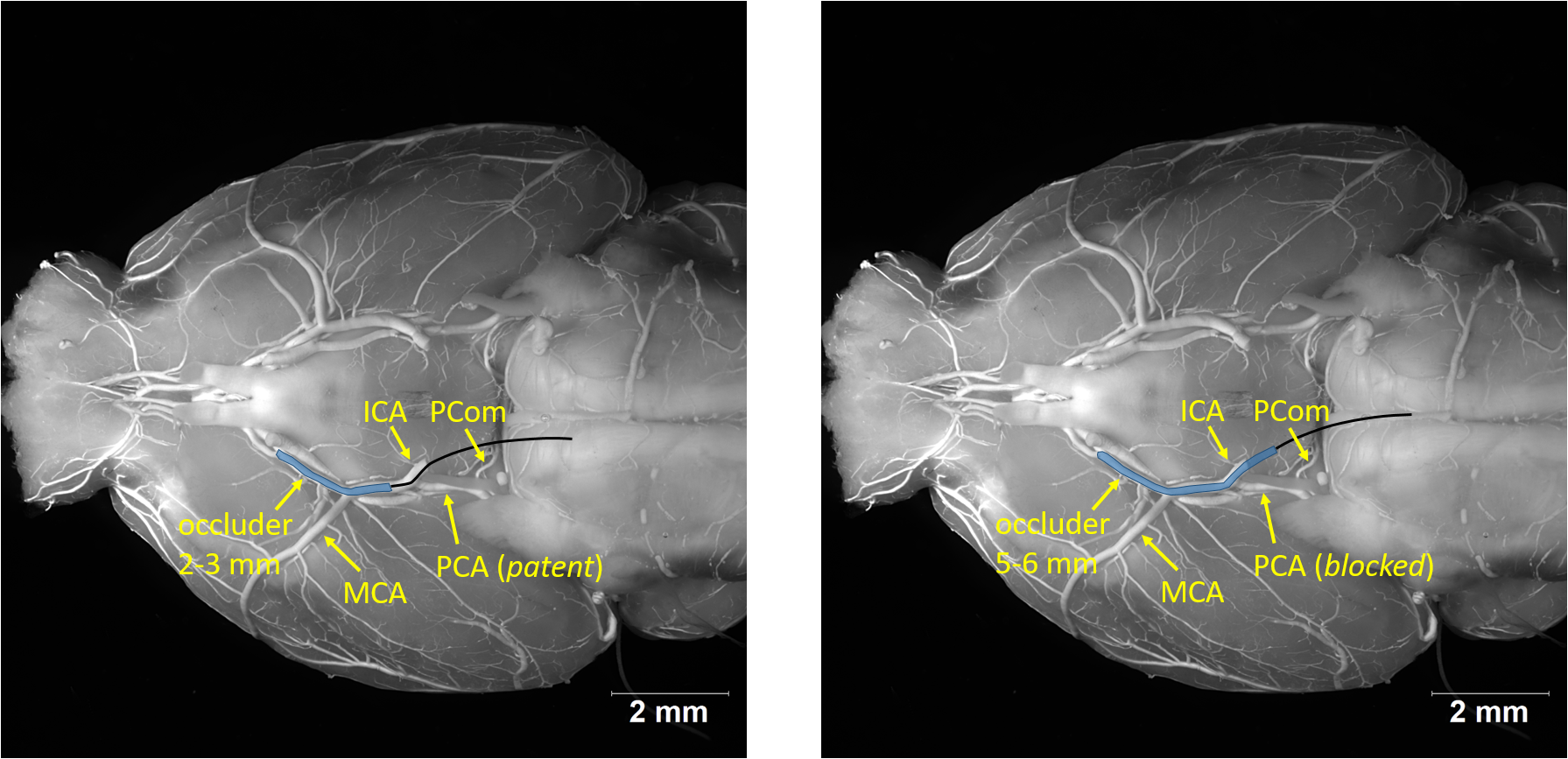

### Supplemental Figure 6

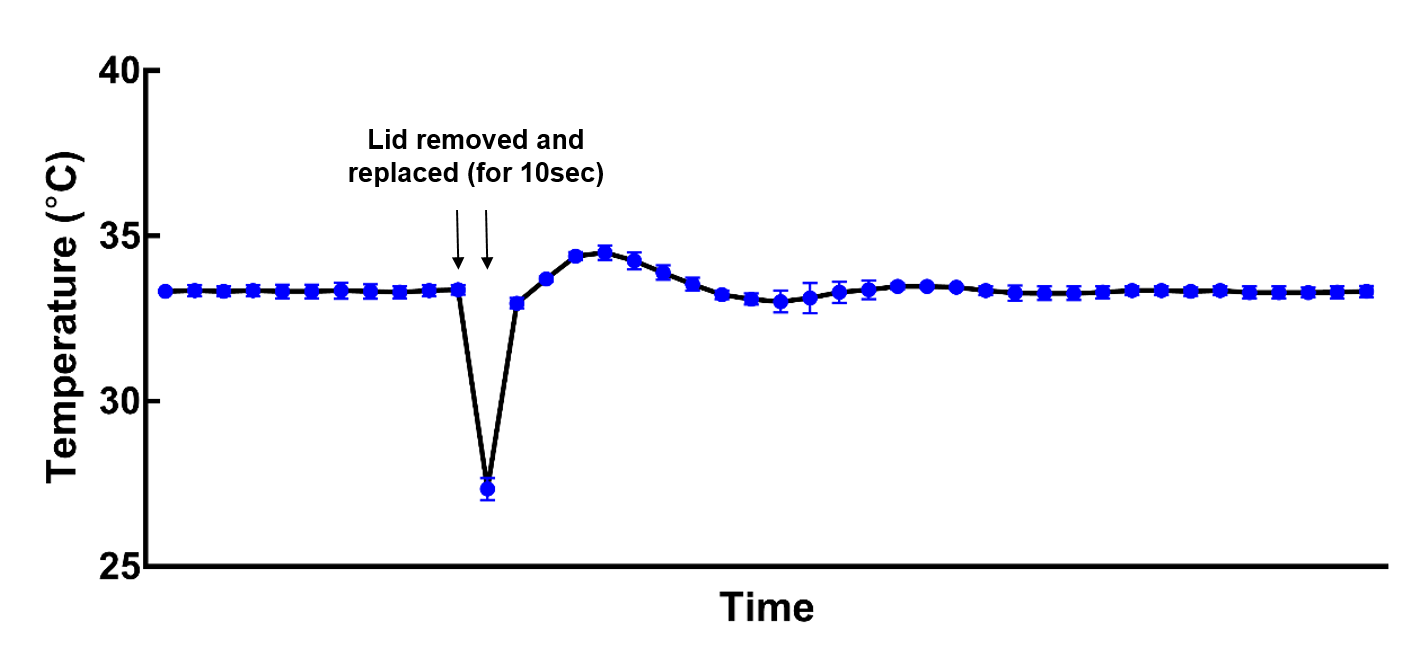
